# Near-Gapless and Haplotype-Resolved *Capsella* Genomes Enable Investigation into Genomic Consequences of Mating System Shifts

**DOI:** 10.64898/2026.07.10.737683

**Authors:** Heng Chen, Robyn A. Emmerson, Rebecca A. Mosher

## Abstract

The shift from outcrossing to self-fertilization is a common evolutionary transition in flowering plants. The genus *Capsella*, comprising the obligate outcrosser *C. grandiflora* and two self-fertile species, *C. rubella* and *C. orientalis*, provides a powerful system to explore genomic consequences of mating system shifts. Despite its utility, existing genomic resources in *Capsella* are fragmented, incomplete, and particularly deficient in repetitive genomic regions, hindering the study of transposable element (TE) dynamics and gene annotation. Here, we present high-quality, chromosome-scale, near-gapless genome assemblies for *C. grandiflora*, *C. rubella*, and *C. orientalis*. Leveraging these improved genomes, we created high-quality genomic resources for *the Capsella* genus by performing comprehensive, *de novo* annotations of protein-coding genes and TEs. Comparative genomic analysis among these species reveals differences in TE abundance, position, and production of small RNAs. These resources provide an unprecedented opportunity to explore how mating system transitions influence genome architecture, TE behavior, and gene evolution. This research also developed a static online platform for *Capsella* genomic resources, Capsella Database (CapBase, www.capsella.uk), to support community use of these resources. Our findings advance understanding of the genomic impacts of selfing and establish a robust foundation for future research into genomics, epigenomics, and evolutionary biology within *Capsella* and related plant systems.

**Significance statement:** The *Capsella* species are model plants for investigating the genomic impacts of the shift from outcrossing to self-fertilization. However, genomic resources for these species remain poor. In this study, we created high-quality genome assemblies and annotations for three *Capsella* species, performed comparative genomic analysis among these genomes, and constructed the first *Capsella* online genomic database, CapBase (www.capsella.uk). These outputs provide a valuable foundation for studying plant genome dynamics following the evolution of self-fertilization.

## Introduction

The shift in mating systems from outcrossing to self-fertilization is one of the most common evolutionary changes in plants (Shimizu & Tsuchimatsu, 2015). This transition is accompanied by a suite of morphological changes, collectively termed the “selfing syndrome”, including reduced pollen production and smaller floral organs (Sicard et al., 2016; Sicard & Lenhard, 2011). Beyond morphology, the shift to self-fertility profoundly affects genomes as well: it reduces heterozygosity, effective population size, and recombination rates, causing accumulation of deleterious alleles (Arunkumar et al., 2015; Barrett et al., 2014; Cutter, 2019; Hough et al., 2013; Lindholm et al., 2016).

Genome size may also be affected by mating-system transitions. Comparative analyses across 205 plant species suggest that outcrossing rate and genome size are positively correlated, consistent with the idea that selfing lineages often evolve smaller genomes (Slotte et al., 2013; Whitney et al., 2010; Wright et al., 2008). One possible contributor to this pattern is transposable element (TE) dynamics; comparisons between selfing and outcrossing *Arabidopsis* species suggest reduced TE abundance in a selfing lineage (Hollister et al., 2011; Lockton & Gaut, 2010). This reduction may reflect both genetic effects, including reduced spread of TEs to new genetic backgrounds, and epigenetic effects, including changes in host-defense mechanisms that control TE silencing (Betancourt et al., 2024; Hollister & Gaut, 2009). However, the expected direction of TE evolution after selfing is not straightforward. In *Arabidopsis lyrata*, for instance, recent population-level work found higher TE copy number and higher mean TE allele frequency in selfing populations than in outcrossers, directly contradicting the expectation of universal TE loss after selfing (Padilla-García et al., 2025). Furthermore, previous studies also suggest that larger genomes can arise in drought/stress environments or at the leading edge of range expansion (Mora-Carrera et al., 2024; Potapenko et al., 2026; Yang et al., 2026), suggesting that mating system might be a minor contributor to genome size variation. Because TEs are also associated with epigenomic phenomena such as genomic imprinting, mating-system transitions may influence not only genome size but also broader epigenomic landscapes (Agren et al., 2014; Brandvain & Haig, 2005, 2018; Slotte et al., 2013; Xiao et al., 2025). Thus, whether the transition to selfing has a predictable impact on genomic and epigenomic landscapes is unclear.

*Capsella*, a small genus of flowering plants in the *Brassicaceae* family, offers an exceptional system for studying the evolution of selfing, as this transition has occurred both recently and repeatedly in *Capsella*. The genus includes three closely related diploid species: one obligate outcrosser, *C. grandiflora*, and two predominantly self-fertile taxa, *C. rubella* and *C. orientalis* (Douglas et al., 2015; Foxe et al., 2009; Sicard & Lenhard, 2018; Sicard et al., 2011). Molecular dating indicates that *C. rubella* split from *C. grandiflora* 50–200 KYA (thousand years ago) when *C. rubella* eliminated its self-incompatibility system, creating a fully self-compatible species (Bachmann et al., 2019; Brandvain et al., 2013; Guo et al., 2009). The origin of selfing in *C. orientalis* is unknown, but might have occurred upon divergence from the *C. grandiflora-C. rubella* precursor, approximately 900 KYA (Douglas et al., 2015; Sicard & Lenhard, 2018).

While *Capsella* has been a powerful model to study the developmental changes associated with mating system transitions (Bachmann et al., 2019; Sicard & Lenhard, 2011; Slotte et al., 2013), analysis of genomic changes in *Capsella* has been hindered by fragmented and incomplete genome assemblies. Most existing genomes were generated from short-read sequencing, resulting in poor coverage of repetitive regions, particularly TEs. This limitation is especially acute for *C. grandiflora*, whose high heterozygosity complicates assembly and scaffolding. As a result, none of the three available *C. grandiflora* assemblies has reached chromosome-level contiguity (Dew-Budd et al., 2024; Dziasek et al., 2024; Slotte et al., 2013). Although chromosome-scale references exist for the other two *Capsella* species (Kasianova et al., 2025; Slotte et al., 2013), numerous gaps remain in these assemblies. This deficiency strictly limits TE-related research, a critical aspect of genome evolution.

Besides genome quality, annotations of protein-coding genes and TEs are other major features of high-quality genomic resources. In the current *Capsella* reference set, gene annotations exist for all three species, but they lack annotation of multiple transcripts (alternative splicing) and untranslated regions (UTRs), and homologous annotation/comparison within the genus. Such deficiencies impede studies of gene evolution, transcriptome dynamics, epigenetic regulation, and molecular biology in *Capsella*. Similarly, the existing TE annotations of *Capsella* assemblies rely either on database-guided methods (*e.g.* RepBase or Dfam) (Bao et al., 2015; Storer et al., 2021) or *de novo* pipelines (*e.g.*, RepeatModeler2 or EDTA) (Flynn et al., 2020; Ou et al., 2019), leading to incomplete and inaccurate TE annotations for the former or inconsistent TE classification for the latter. Considering that TEs may be significantly influenced by the selfing transition, an accurate and well-classified TE annotation is obligatory.

To enable the study of *Capsella* genomes, we generated near-gapless, chromosome-scale genomes for *C. grandiflora*, *C. rubella*, and *C. orientalis* and annotated genes and TEs *de novo* in each genome. Here, we present a comparative analysis of the genome, gene, and TE levels among these *Capsella* species. The results suggest that while protein-coding genes are highly conserved within the *Capsella* genus, TE content is substantially different in each species, perhaps due to enhanced sRNA-mediated host defense in inbreeders. Non-TE repeats, especially in the centromere region, also contribute to the greater genome size in *C. grandiflora*. The resulting high-quality genomic resources will facilitate future research into genomics, epigenomics, and evolutionary biology within *Capsella*, deepening our understanding of how mating-system transitions shape plant genomes.

## Results

### Assembly of near-gapless *Capsella* genomes

The three *Capsella* genomes were sequenced with high-fidelity long reads and chromatin conformation capture sequencing. Each genome was assembled into eight nearly gapless chromosomes that were substantially longer and more contiguous than previously available genome assemblies (**Table 1 and Supplementary Figure 1**). Compared to the public *C. rubella* and *C. orientalis* chromosome-level assembly (Kasianova et al., 2025; Slotte et al., 2013), our assembly was 24% longer, especially in repetitive regions. Improvements in the *C. grandiflora* genome were even more pronounced, as the genome was not previously scaffolded into pseudomolecules (**Table 1 and Supplementary Table 1**).

**Table 1.** *Capsella* genome assembly statistics.

| Species/Name<sup>a</sup> | Assembly level | Length (Mbp) | No. scaffolds | No. gap<sup>b</sup> | Scaffold N50 (Mbp) | Scaffold L50 |
| --- | --- | --- | --- | --- | --- | --- |
| *Capsella rubella* |  |  |  |  |  |  |
| Capsella rubella v1.1 | Chr level | 135 | 853 | 23,427 | 15.1 | 4 |
| **Crubella_RAM_v1.1** | **Near gapless** | **168** | **550** | **69** | **20.4** | **4** |
| *Capsella grandiflora* |  |  |  |  |  |  |
| Capsella grandiflora v1.1 | Contig level | 105 | 4,997 | N/A | 0.1 | 223 |
| UAZ_Capgrnd_2 | Contig level | 145 | 577 | N/A | 2.3 | 17 |
| Cgrandiflora_new | Contig level | 180 | 537 | N/A | 0.7 | 49 |
| **Cgrandiflora_hapA_RAM_v1.1** | **Near gapless** | **278** | **620** | **37** | **24.4** | **5** |
| **Cgrandiflora_hapB_RAM_v1.1** | **Near gapless** | **231** | **1995** | **36** | **14.8** | **7** |
| *Capsella orientalis* |  |  |  |  |  |  |
| CapOri_SW_1.1 | Contig level | 111 | 11,478 | N/A | 0.2 | 1758 |
| Corientalis_genome | Near gapless | 131 | 63 | 22 | 16.2 | 4 |
| **Corientalis_RAM_v1.1** | **Near gapless** | **185** | **87** | **25** | **22.0** | **4** |

<sup>a</sup> **Species/Name:** The name of the species and assemblies. Genomes sequenced and assembled by the current research are highlighted in bold. The official ID of released assemblies can be found in Table S1.
<sup>b</sup> **No. gap:** The number of gaps in the pseudochromosomes. ≥15 contiguous Ns is determined as one gap.

Our three *Capsella* genomes were highly syntenic (**Figure 1A** and **Supplementary Figure 2**), showing variability primarily in the centromeric satellite repeats. One notable break in synteny between our *C. rubella* and the public *C. rubella* assembly was an approximately 2 Mb inversion on the short arm of chromosome 7 (**Figure 1B, Supplementary Figure 1**). The inversion breakpoint was in the middle of a 4Mb contig in our assembly, giving us high confidence in the structure of this region. The breakpoint coincided with a 30 kb gap in the public assembly, suggesting that scaffolding might have inadvertently flipped the end of the chromosome.

**Figure 1.**
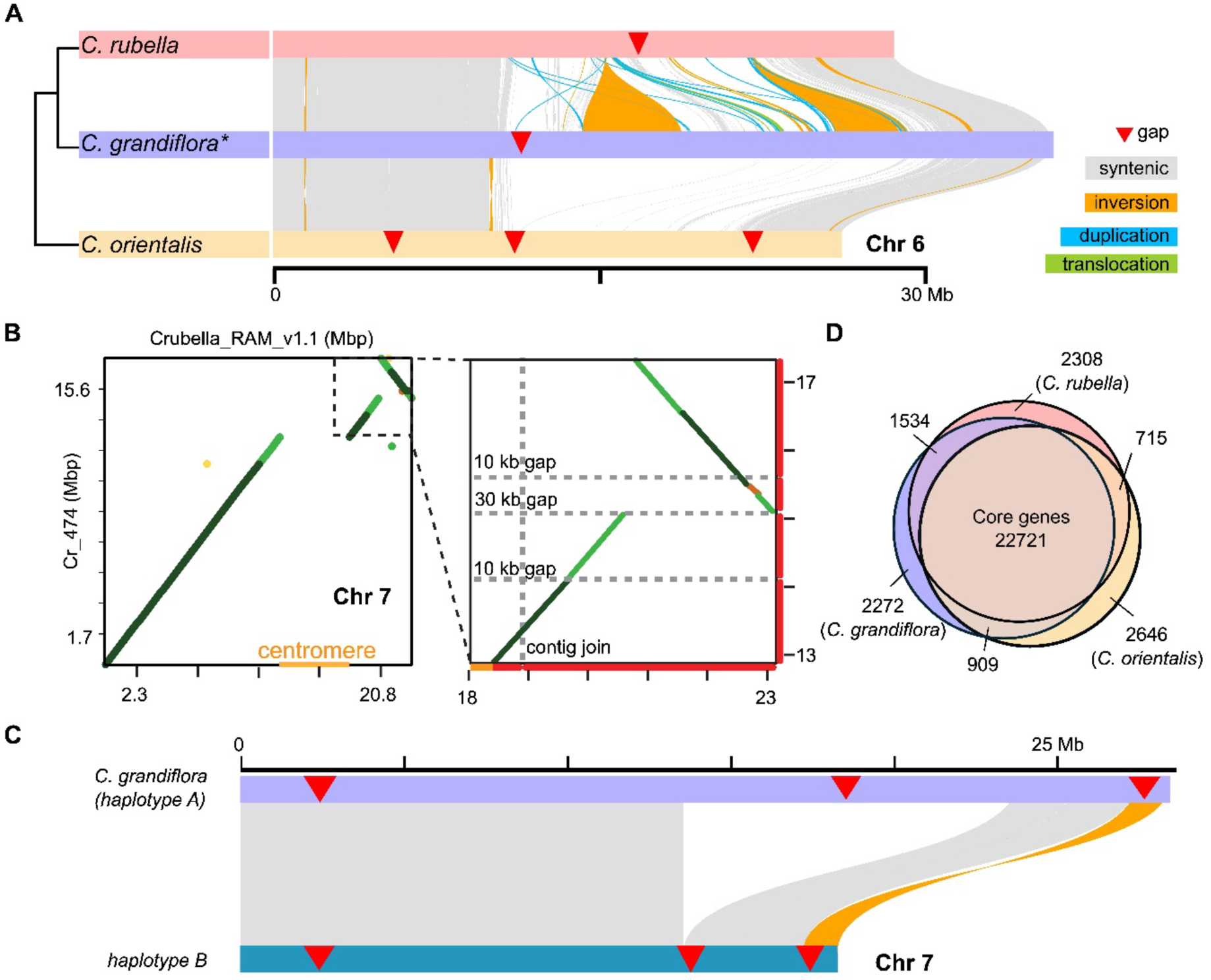
Synteny among *Capsella* genomes. **(A)** k-*mer* based analysis shows extensive synteny in chromosome arms. All chromosomes, including gene-based synteny dot plots, can be found in Supplementary Figure 1. **(B)** Synteny plots demonstrate an approximately 2 Mb inversion at the end of chromosome 6 in our assembly relative to the published *C. rubella* genome. Scaffolding gaps are shown to identify contig joins. **(C)** Venn diagram depicting orthologous genes (syntenic and non-syntenic) shared by *Capsella* genomes. The 22,721 core genes were defined based on *C. rubella*. The number slightly varies in *C. grandiflora* (22,804) and *C. orientalis* (22,672), due to species-specific paralogs. **(D)** Riparian plot showing extensive synteny in the chromosome arms between *C. grandiflora* haplotypes (chromosome 7 shown). The synteny plots for all chromosomes can be found in Supplementary Figure 3.

In addition to the nuclear genome, complete chloroplast and mitochondrial genomes (154-155 kb and 287-288 kb, respectively) were obtained from each of the three *Capsella* species. Long read sequencing allowed us to identify two versions of the chloroplast genome in each species due to a 26 kb inversion. Similarly, extensive rearrangements in the mitochondrial genomes generated 4-22 confirmed structures in each species. A single version of each chloroplast and mitochondrial genome is included in our genome assemblies; all assemblies can be found in **Supplementary File 1**.

As an outbreeding species, *C. grandiflora* is largely heterozygous, and we therefore assembled two haplotypes (**Figure 1C**, **Table 1**): haplotype A (hapA, the reference assembly discussed above) and haplotype B (hapB, the alternative assembly). These two assemblies each formed 8 chromosomes, and were highly syntenic (**Figure 1C and Supplementary Figure 3A-B**). We used hapA as the representative assembly because hapA is longer, has fewer small scaffolds, and has better N50 and L50 scores (**Table 1**). In addition to many small duplications and translocations, there are some larger differences between haplotypes at the genomic level, including small inversions in chromosomes 1, 5, and 7 (**Supplementary Figure 3A**). Some of these inversions correspond to breakpoints in the assembly, suggesting they might represent errors in assembly rather than chromosomal polymorphisms.

### High-quality *de novo* genome annotation of *Capsella*

Protein-coding genes for each genome were annotated by a custom pipeline integrating *ab initio* prediction, RNA seq- and protein-based annotation, and homologous reference-based curation. Non-coding genes were also defined by various pipelines. As expected, the overall number of annotated genes and transcripts (approximately 27,000 and 60,000, respectively) is similar among these species (**Table 2**). The high completeness and consistent lineage scores from OMArk, a proteome and gene family-based tool for gene annotation quality control (Nevers et al., 2025), indicate that our annotation is reliable and complete (**Supplementary Table 2**).

**Table 2.** *Capsella* genome annotation statistics.

| Genome Name | Protein-Coding Genes | Transcripts | ncRNA<sup>a</sup> | lncRNA<sup>b</sup> |
| --- | --- | --- | --- | --- |
| Crubella_RAM_v1.1 | 27,455 | 57,937 | 7,161 | 3,950 |
| Corientalis_RAM_v1.1 | 27,107 | 63,446 | 4,973 | 3,552 |
| Cgrandiflora_RAM_v1.1_hapA | 28,388 | 60,451 | 13,994 | 4,257 |
| Cgrandiflora_RAM_v1.1_hapB | 28,087 | 60,350 | N/A | N/A |

<sup>a</sup> ncRNA includes rRNA, tRNA, snRNA, and similar small non-coding RNA.
<sup>b</sup> lncRNA includes non-protein-coding transcripts longer than 200nt.

We next compared protein-coding genes in the three *Capsella* genomes with a custom pipeline that combined sequence identity and synteny (**Figure 1D**, **Supplementary Table 3 - 5)**. Approximately 83% of genes (22,721) are present in all three genomes and were defined as “core” genes. Most core genes were found on the 8 chromosomes, where they were evenly distributed. There are more (1534) *C. rubella* - *C. grandiflora* homologs (*i.e.,* no homologs in *C. orientalis*) than *C. rubella* - *C. orientalis* and *C. grandiflora* - *C. orientalis* homologs. This is not surprising given the more recent divergence between *C. rubella* and *C. grandiflora* (Slotte et al., 2013). In contrast to gene content, the number of short non-coding small RNAs (*e.g.*, tRNAs, snoRNAs) varies substantially between *Capsella* species (**Table 2**), primarily due to differences in the size of rRNA and tRNA repeat arrays in the assembly.

We also performed gene annotation for *C. grandiflora* hapB. Similar to hapA, we annotated about 28,000 genes with 60,000 transcripts (**Table 2**). About 94% of these genes (∼26,400) are conserved between haplotypes (**Supplementary Figure 3C and Supplementary Table 5**), suggesting a closer relationship between the two haplotypes than between *C. grandiflora* and *C*. *rubella*.

Together, these results demonstrate strong conservation of protein coding genes between *Capsella* species, with differences scaling with time since divergence. The supplied tables of syntenic homologs (**Supplementary Table 3 - 5**) will be a useful tool for researchers working in the genus. Moreover, this research also developed the Capsella Database (CapBase, www.capella.org.uk), a static online platform that integrates assemblies, annotations, and related bioinformatic tools (an ortholog finder and a genome browser), to support community use of these resources.

### Variable TE accumulation in *Capsella*

To understand TEs, the most variable part of a genome, we first developed species-specific repeat libraries with customized pipelines and then annotated each genome with EDTA (Ou et al., 2019) (**Table 3**, **Supplementary Table 6**). The total TE content in *C. grandiflora* is higher than *C. rubella* and *C. orientalis*, whether measured as number of annotations, absolute TE length, or proportion of the genome (**Table 3**, **Figure 2A, Supplementary Figure 4**). This observation contrasts with previous analysis, which detected a slightly higher TE copy number in *C. rubella* compared to *C. grandiflora* and sharply lower numbers of TEs in *C. orientalis* (Agren et al., 2014). Higher TE accumulation in the outbreeder *C. grandiflora* is consistent with TE dynamics following the transition to selfing in *Arabidopsis thaliana* and *Arabidopsis lyrata* (Hollister et al., 2011). Similar TE load in *C. orientalis* and *C. rubella* also suggests there may be a predictable influence on TE dynamics following the transition to selfing. However, closer analysis indicates that evolution may have different impacts on individual TE subclasses. Class II elements, especially TIR elements, are less common in *C. grandiflora* relative to the other *Capsella* species (**Figure 2A**, **Supplementary Figure 4**).

**Figure 2.**
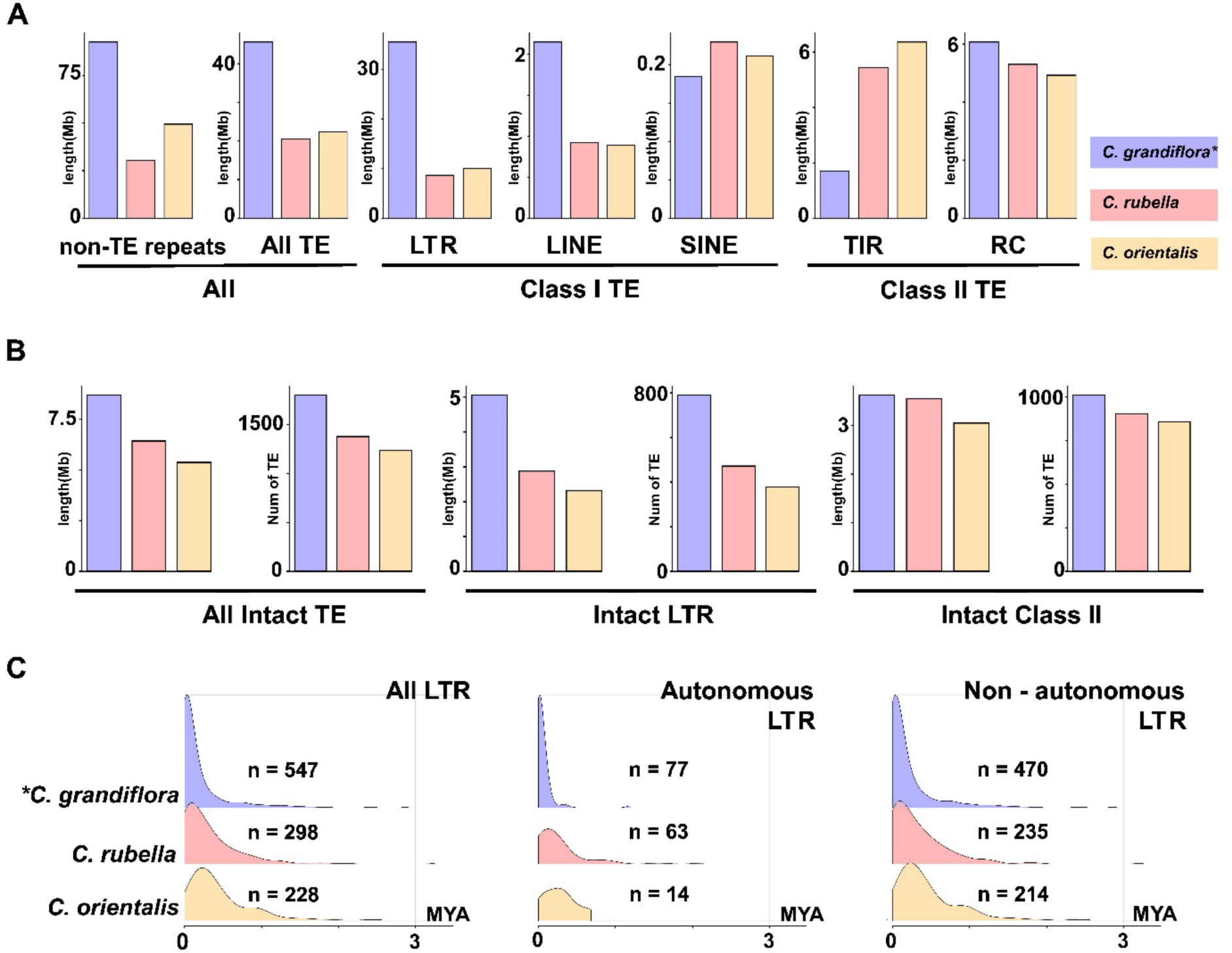
Different intact TE abundance implies more recent TE transposition in *C. grandiflora*. **(A)** TE abundance in *Capsella* genomes. The non-TE repeats include Satellite DNA, Simple repeats, Tandem repeats, rRNA, tRNA, and Unknown repeats. The TIR includes TIR, MITE, and nMITE elements. The purple, pink, and yellow bars are *C. grandiflora*, *C. rubella,* and *C. orientalis*, respectively. **(B)** Intact TE abundance in *Capsella* genomes. EDTA defined the intact TEs. **(C)** LTR insertion time predicted by MegaLTR. The X-axis is the LTR insertion time in MYA (Million Years Ago), and the Y-axis is the number of inserted LTRs.

**Table 3.** Count of *Capsella* TEs.

| TE_Type | Crubella_RAM_v1 | Corientalis_RAM_v1 | Cgrandiflora_RA<br>M_v1_hapA | Cgrandiflora_R<br>AM_v1_hapB |
| --- | --- | --- | --- | --- |
| **Class I TE** | 9,702 | 8,359 | 16,433 | 14,304 |
| **Class II TE** | 14,751 | 17,550 | 10,585 | 11,279 |
| **All TE** | 23,110 | 27,252 | 27,018 | 25,583 |
| **Non TE repeats <sup>a</sup>** | 9,889 | 13,416 | 30,406 | 16,821 |

<sup>a</sup> **Non TE repeats** include tRNA, rRNA, tandem repeats, simple repeats, satellite DNA, and unknown repeats.

The length per TE is substantially higher in *C. grandiflora*, implying that *C. grandiflora* may contain longer and/or more intact TEs. We therefore investigated the abundance of intact TEs in *Capsella* species. *C. grandiflora* shows the highest intact TE content and the largest number of intact TEs, particularly LTRs, compared to the other two species (**Figure 2B**). Because insertions are expected to degrade and become smaller over time through partial deletions, we measured the age of LTRs elements and observed younger insertion ages for LTR elements in *C. grandiflora* (**Figure 2C**), The reduced number of intact elements in *C. orientalis* and *C. rubella*, along with their older age profile, implies fewer recent transpositional events, indicating that reduction in transposition activity might be a consequence of the transition to selfing.

We next assessed the density of TEs in the genome and across chromosome arms in each species. For most elements TE density (percentage of the genome annotated as TE) is similar in *C. rubella* and *C. orientalis,* but different in *C. grandiflora* (**Supplementary Figure 4**), which might indicate a common trajectory for TEs following the selfing transition. When viewed in the context of chromosomes, both Class I and Class II elements were more frequent in chromosome arms of all three genomes. However, there is an enrichment of Class I TEs in *C. grandiflora* centromeres and a depletion of these elements in *C. orientalis* centromeres (**Figure 3A, B).**

**Figure 3.**
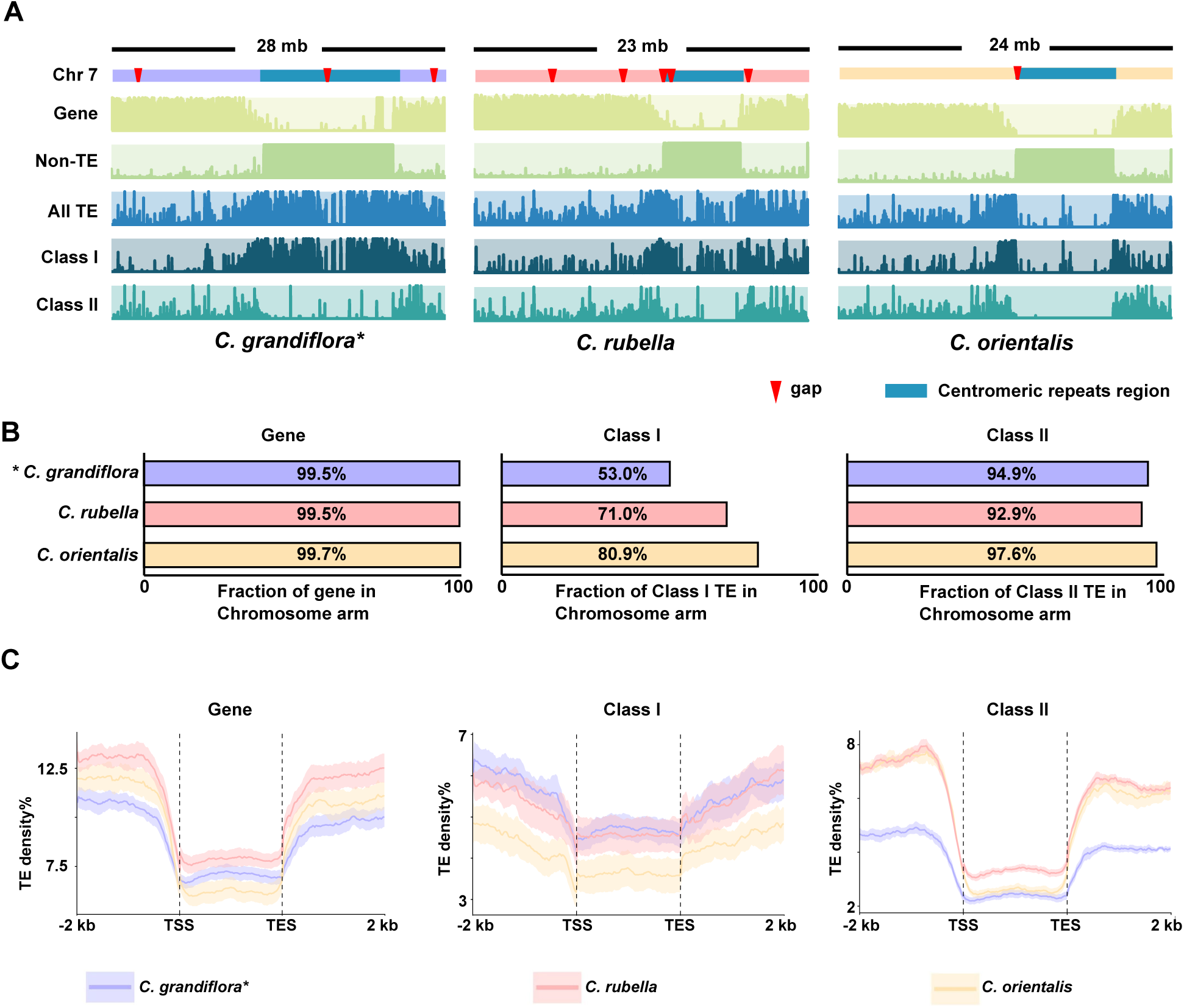
Class 1 TEs are enriched in chromosome arms in selfing *Capsella* species. **(A)** Abundance of different genomic features in the Chr 7 of *Capsellas*. Red triangles mark gaps in the assembly. The centromeric repeat-enriched region is highlighted with blue bars. The non-TE repeats include Satellite DNAs, Simple repeats, Tandem repeats, rRNAs, tRNAs, and Unknown repeats. Class I TE includes LTRs, SINEs, and LINEs. Class II TE includes Helitrons, TIRs, MITEs, and nMITEs. **(B)** The bar plots show the percentage of total length overlapped by annotation of genes, Class I TEs, and Class II TEs in chromosome arms. **(C)** Metagene plots show TE density in the gene body (transcription start site (TSS) – transcription end site (TES)) and 2 kb flanking regions. The error bars show standard errors calculated across the eight chromosomes.

To further investigate the distribution of TEs relative to genes, we plotted TE density across genes and their flanking regions (**Figure 3C**). Firstly, TE density is higher in gene-flanking regions than in gene bodies. This pattern is expected due to selection against insertions that disrupt gene sequence. Moreover, the difference in TE density between gene bodies and flanking regions is more pronounced for Class II elements, indicating that the capacity of different types of TE to insert or be maintained in gene bodies is varied in *Capsella*. Distribution of Class I elements in and around genes is very similar in *C. rubella* and *C. grandiflora* (**Figure 3C**), despite much higher genome density of Class I elements in *C. grandiflora* (**Supplementary Figure 4**). This pattern likely arises due to the abundance of Class I elements in *C. grandiflora* centromeres and suggests that although there are more active Class I elements in *C. grandiflora*, there is little difference in the impact on protein coding genes. Distribution of Class II elements relative to genes largely follows their density in the genomes.

In brief, we observed differences in TE composition and abundance among the three *Capsella* species. These differences support the hypothesis that TE dynamics, especially LTR dynamics, dampened following speciation events for *C. rubella* and *C. orientalis*, and suggest that there might be a common trajectory for TEs upon the evolution of selfing. Moreover, the distribution of different types of TEs across chromosomes and relative to protein-coding genes are different, indicating mating system transition might show different impacts on determining fate of DNA based TEs and retrotransposons.

### Transposable element silencing may differ between breeding strategies

The primary mechanism suppressing TE mobility is epigenetic silencing, including DNA methylation induced by 24-nt siRNAs (Matzke & Mosher, 2014). To understand whether host defense against TEs quantitatively differs depending on mating system, we mapped sRNA-seq data from leaves of each *Capsella* species. When compared to the publicly-available genome assemblies (Kasianova et al., 2025; Slotte et al., 2013), more sRNAs mapped to our genomes (**Figure 4A, and Supplementary Table 7 - 8**), further highlighting the improved nature of our assemblies. Our *C. grandiflora* assembly more than doubled the number of mapped reads compared to the publicly available assembly, particularly for the 23-24-nt size class. *C. rubella* and *C. orientalis* saw modest increases of ∼3% and ∼0.8% increase, respectively, with 23-24-nt sizes leading this difference. While improvements in sRNA mapping in *C. orientalis* could be due to mapping sRNAs to the same genotype that was sequenced, for *C. rubella*, both the public genome and sRNA data come from the same accession.

**Figure 4.**
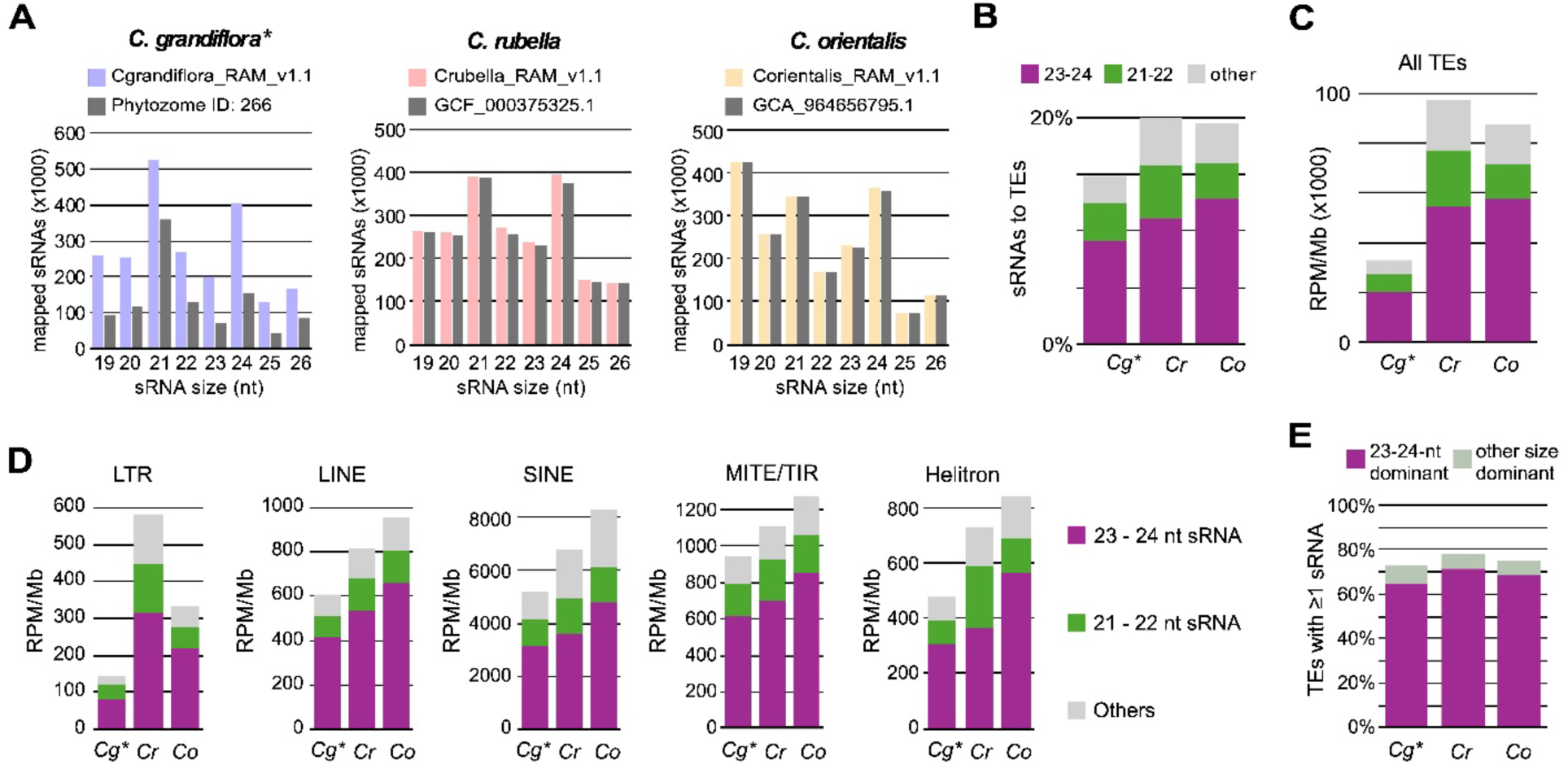
Small RNA benchmarking and alignment over TEs. **(A)** Size profiles of small RNAs (sRNA) that map to our new assemblies (blue, pink, yellow) and the current publicly available assemblies (grey) by species. **(B)** Fraction of sRNA libraries mapping to TEs in each genome, by size class. **(C)** sRNA mapping to TEs adjusted by Mb of TE in each genome and library size. **(D)** As in C, but broken down by TE subclasses. **(E)** The percentage of TEs producing sRNAs in each species. 23-24 nt dominant TEs produce more 23-24 nt sRNAs than 21-22 nt sRNA, or irregular sizes (19, 20, 25, and 26 nt) of sRNA. The other size dominant TE means that the most abundant sRNA produced by the TE are 21-22 nt or irregular sizes.

We then investigated whether sRNA mapping to TEs differed between *Capsella* species. 15% of 19-26-nt sRNAs overlapped TE sequences in *C. grandiflora*, while approximately 20% of sRNAs matched to TEs in *C. rubella* and *C. orientalis* (**Figure 4B, and Supplementary Table 7 - 8**). As expected, the majority of TE-aligning sRNAs are 23-24-nt for each species. The decreased percentage of sRNAs mapping to TEs in *C. grandiflora* is despite the increased TE content in this species. When read mapping is adjusted for total length of TEs in each genome, there is a sharp reduction in sRNAs from TEs in *C. grandiflora* (**Figure 4C**). When analyzed by class and type of TE, there is consistently lower sRNA RPM per Mb TE in *C. grandiflora* and the highest RPM/Mb in *C. orientalis*, with the exception of LTR elements where *C. rubella* is highest (**Figure 4D**).

The increase in sRNA density across TEs in *C. rubella* and *C. orientalis* relative to *C. grandiflora* could result from greater sRNA production from TEs that are recognized by the sRNA defense pathway or from a larger number of TEs being recognized and producing sRNAs. To distinguish these options, we measured the number of TEs producing at least one sRNA. A slightly greater percentage of TEs produced sRNAs in *C. orientalis* and *C. rubella* than *C. grandiflora* (**Figure 4E**), however this difference cannot fully explain the 2.5-3-fold increase in sRNA production (RPM/Mb) observed in these species (**Figure 4C-D**). Together, these observations suggest that after a selfing transition, *Capsellas* have stronger sRNA-based TE defense, which might explain the decreased number of TEs in inbreeding *Capsellas*.

## Discussion

We assembled near-gapless, chromosome-scale genomes and produced high-quality gene and TE annotations for *C. grandiflora*, *C. rubella*, and *C. orientalis*. These resources provide a robust foundation to interrogate how the transition to self-fertilization reshapes genomes, including TE defense. The results indicate that genes are highly conserved within the genus, while TEs, especially LTR retroTEs, are suppressed following the transition to selfing, possibly due to enhanced sRNA-mediated silencing.

Selfing is predicted to reshape TE dynamics, however the precise fate of TEs following a transition to selfing is unclear (D. Charlesworth & Wright, 2001). Reduced outcrossing limits the spread of a TE through a population, suggesting lower TE content in inbreeders (Boutin et al., 2012). Reduced recombination efficiency due to increased homozygosity will also maintain linkage between potentially active TEs and loci responsible for TE silencing (*e.g.*, complex TE insertions that produce double-stranded RNA and trigger siRNA formation), suggesting stronger suppression of TE mobility in inbreeding species (B. Charlesworth & Langley, 1986). However, reduced recombination will also slow the removal of TEs and reduce the efficacy of selection against mildly deleterious elements (D. Charlesworth & Wright, 2001). These pressures would increase TE abundance in inbreeders. Empirical data, including at population-scale, is necessary to test these theoretical assumptions and understand the role of mating system transition on TE dynamics (Agren et al., 2014). However, TEs are among the most difficult portions of a genome to assemble, which has limited efforts to procure empirical evidence. Our near-gapless, chromosome-scale genomes include highly assembled centromeres and pericentromeres, recover long intact elements, and anchor TE annotations to a consistent gene and synteny framework. Furthermore, species-specific repeat libraries and a unified TE curation eliminate cross-species classification bias, allowing “apples-to-apples” contrasts of TE load, intactness, age, and spatial patterning relative to genes and heterochromatin.

Previous analysis suggested a similar copy number of each major type of TEs in *C. grandiflora* and *C. rubella*, with sharply lower TE numbers in *C. orientalis* (Agren et al., 2014). However, this data was based on mapping paired-end short read data to a *C. rubella* reference genome and likely suffered from reference genome bias. In our genome assemblies the outbreeder *C. grandiflora* harbors substantially higher numbers of Class I (RNA-based) elements, but has sharply reduced levels of Class II (DNA-based) elements relative to the inbreeding species *C. rubella* and *C. orientalis.* This might suggest that RNA- and DNA-based TEs have different dynamics following the transition to selfing.

Our evidence suggests that these differences in TE populations derive in part from suppression of TE activity following the transition to selfing. Firstly, overall, we detect more LTR and LINE elements, more intact LTRs, and younger LTRs in the outbreeder *C. grandiflora*, than in either of the inbreeding species, *C. rubella* or *C. orientalis* (Figure 2). This pattern suggests ongoing retrotransposition in *C. grandiflora* and suppression of LTR activity in the inbreeding species. Our observations in *Capsella* are consistent with increased TE accumulation and younger TE insertions in the outbreeder *Arabidopsis lyrata* compared to the inbreeder *A. thaliana* (Slotte et al., 2013). Secondly, we detect a higher load of TE-derived siRNAs in the inbreeding species, especially those matching LTR elements (Figure 4), indicating that host defenses are more active following selfing. Similarly, a higher proportion of TEs match siRNAs in *A. lyrata* compared to *A. thaliana* (Hollister et al., 2011), suggesting that increased TE defense might be a common outcome following the transition to selfing. However, it is important to recognize that TE dynamics described here might be a consequence of the low effective population size upon breakdown of self-incompatibility rather than mating system *per se* (Mora-Carrera et al., 2024; Yang et al., 2026).

In addition to more complete coverage of TEs, the *Capsella* genomes described here include nearly complete centromeres. Although larger genomes are frequently attributed to the accumulation of TEs, more than half of the size difference between *C. grandiflora* and its inbreeding relatives is due to larger centromeric repeat arrays. In addition, the *C. grandiflora* centromeric arrays are enriched for LTR retrotransposons, further increasing genome size versus the selfing species. Whether elimination of DNA in low-recombination, repeat-rich regions such as centromeres is a feature of selfing species is an open question that will require additional genome assemblies like those created here.

## Materials and Methods

### Genetic material

*C. grandiflora* (accession 83.17) and *C. rubella* (Monte Gargano) were a gift from Dr. Mark Beilstein (University of Arizona); *C. orientalis* (accession Co1983) was donated by Dr. Michael Lenhard (University of Potsdam) (Woźniak et al., 2020). All plants were grown in growth chambers at 20 ℃ and 50% relative humidity with 16 h of daylight. To construct the PacBio and Omni-C (HiC) sequencing library, fresh leaves of each species were collected and flash-frozen in liquid nitrogen before shipping to Cantata Biosciences (CA, USA). For RNA sequencing, unfertilized ovules or leaves (three biological replicates each) of *C. rubella* and *C. grandiflora* were collected. Total nucleic acids were extracted as described previously (White & Kaper, 1989) and sequenced at SeqCenter (PA, USA).

### Genome Sequencing and Assembly

PacBio Circular Consensus Sequencing (CCS) and Illumina Omni-C libraries were prepared and sequenced by Cantata Biosciences (CA, USA). Raw sequence coverage of PacBio libraries was 453x (*C. grandiflora*), 315x (*C. rubella*) and 208x (*C. orientalis*); Omni-C libraries were approximately 30x for each genome.

First, draft genomes were *de novo* assembled from PacBio HiFi reads using Hifiasm with default parameters (H. Cheng et al., 2022). Blast results of the Hifiasm output assembly against the NT database were used as input for Blobtools2 and scaffolds identified as possible contamination were removed from the assembly. Finally, purge_dups3 was used to remove haplotigs and contig overlaps (Altschul et al., 1990; Guan et al., 2020; Laetsch & Blaxter, 2017).

Next, the *de novo* assembly and OmniC library reads were used to scaffold the contigs with HiRise (Putnam et al., 2016). Because of the low number of contigs in the draft assembly, all scaffolding joins were manually confirmed and corrected by comparison to the public *C. rubella* genome (Phytozome genome ID: 474)(Goodstein et al., 2012; Slotte et al., 2013).

### Organelle Genome Assembly and Annotation

To assemble the *Capsella* chloroplast genomes *de novo*, GetOrganelle was used with recommended parameters for PacBio HiFi reads and reference to public *Capsella* chloroplast assembly from NCBI (NC_027693.1, NC_028517.1 and NC_081111.1) (Jin et al., 2020; Wu, 2016; Wu & Ma, 2016). To assemble the mitochondrial genomes *de novo*, PMAT2, GetOrganelle, and Bandage were used (Bi et al., 2024; Jin et al., 2020). Finally, complete organelle genomes were merged with the whole genome assemblies, and small contigs that perfectly matched the organelle genomes were removed. The GeSeq was used to annotate the chloroplast genome, while the LiftOn was used with reference to the Araport11 for mitochondrial genome annotation (Chao et al., 2025; C.-Y. Cheng et al., 2017; Tillich et al., 2017).

### Annotation of Protein Coding Genes

First, each genome assembly was soft-masked by RepeatMasker-4.1.6 with a *de novo* repeat library constructed by RepeatModeler-2.0.5 (Flynn et al., 2020; Tarailo-Graovac & Chen, 2009). The soft-masked genomes were then annotated with BRAKER3 pipeline, which integrated *ab initio* prediction, RNA-seq-based annotation, and protein-based (*Brassicales* excluded OrthoDB v11 protein sets) annotation (Gabriel et al., 2024; Kuznetsov et al., 2023). In addition to publically-available RNA-seq data, additional RNA-seq libraries from ovules and leaves were included (**Supplementary Table 9**). The flag --busco_lineage eukaryota_odb10 was employed to enhance the accuracy of the final gene set by minimizing missing BUSCOs.

Next, draft annotations were refined by incorporating UTRs and alternative splicing variants identified through the PASA pipeline (Haas et al., 2003), which used unmasked genomes and a comprehensive transcriptome database. This database combined Trinity *de novo* RNA-seq assemblies, Trinity genome-guided RNA-seq assemblies, and StringTie2 transcript structures (Grabherr et al., 2011; Kovaka et al., 2019). The RNA-seq data for each of these was the same as the above (**Supplementary Table 9**). Five cycles of PASA annotation updates were performed for each genome to maximize the incorporation of transcript alignments into gene structures.

Finally, the improved annotations were further refined and polished by a customized pipeline. We first correct mis-merged genes, the genes whose isoforms form two or more distinct clusters of transcripts that do not share any overlapping genomic coordinates. Then, we corrected abnormal genes and transcripts. Transcripts were deemed abnormal if they satisfied both of the following conditions: 1) they were shorter than 200 bp (tiny mRNAs) or their parental genes exceeded 10 kb in length (giant genes); and 2) they lacked homologs in the UniProt database (UniProt Consortium, 2023) and did not have homologs in Araport11 (*Arabidopsis thaliana*) (C.-Y. Cheng et al., 2017), Chiifu v4.0 (*Brassica rapa*) (L. Zhang et al., 2023), Capsella rubella v1.1 (*Capsella rubella*) (Slotte et al., 2013), or the other two genomes in this research. Abnormal transcripts were removed, and giant genes were reannotated by Helixer with a pre-trained model, land_plant_v0.3_a_0080.h5 (Stiehler et al., 2021; UniProt Consortium, 2023). OMArk was utilized to assess the completeness and the consistency of the gene repertoire, rather than BUSCO because a BUSCO lineage was specified to execute the BRAKER3 pipeline, which could introduce bias in the estimation (Nevers et al., 2025).

### *Capsella* Genus Core Genes Definition

The homologous genes of three *Capsella* species were defined based on sequence similarity-and synteny-based information. In brief, the candidate homologous genes were first identified by DIAMOND (Buchfink et al., 2015) with a stringent threshold of 95% sequence identity. Then, the syntenic homologies were identified by comparing reference-based annotations (generated by LiftOn (Chao et al., 2025)) and *de novo* annotations. Finally, a custom script was employed to determine *Capsella* core genes by integrating sequence identity- and synteny-based candidate homologs (see **Data and Code Availability**).

### Annotation of non-coding Genes

Various pipelines were used to annotate non-coding RNAs (ncRNAs). Ribosomal RNAs (rRNAs) and transfer RNAs (tRNA) were annotated by RNAmmer and tRNAscan-SE, respectively (Chan et al., 2021; Lagesen et al., 2007), while the other conserved ncRNAs (e.g. small nuclear RNAs (snRNAs)) were annotated by Infernal with Rfam database (Kalvari et al., 2018; Nawrocki et al., 2009). Long non-coding RNAs (lncRNAs) were annotated by FEELnc with RNA-seq data (StringTie2 assembled transcripts) (Wucher et al., 2017). All ncRNA annotations for a species were then integrated into a unified generic feature format version 3 (gff3) file.

### Transposable Elements and Centromeric Regions Identification

To annotate TEs, the Extensive *De novo* TE Annotator (EDTA) and customized *Capsella* TE libraries were employed (Ou et al., 2019). We first *de novo* constructed a TE library for each genome based on the approach 3 of a TE annotation protocol (Benson et al., 2025). The library was then trimmed by TEtrimmer and further classified by TEsorter and DeepTE (Qian et al., 2025; Yan et al., 2020; R.-G. Zhang et al., 2022). Finally, the TEs for each species were annotated by EDTA with their own TE library. Intact TEs were identified by EDTA, and TE ages were estimated with MegaLTR (Mokhtar & El Allali, 2023). Annotations of 5S Short Interspersed Nuclear Element (SINE/5S, SINE derived from 5S rRNA) in *C. rubella* and *C. grandiflora* (hapA) were manually marked as rRNA, non-TE repeats, if they were too long to be a SINE (typically 100 and 700 bp in length (Huang et al., 2012)) and overlapped with an rRNA tandem array (more than 5 rRNA annotations).

To determine the candidate centromeric regions, customized scripts (see **Data and Code Availability**) were used based on local enrichment of non-TE repeats (including satellite DNA, tandem repeats, simple repeats, etc.), together with depletion of gene annotation. Each pseudochromosome was divided into 20 kb windows and windows with ≥40% non-TE repeats and ≤15% geneswere merged if they were within 200 kb. Merged regions shorter than 200 kb were removed and remaining merged regions defined the centromeres.

### Small RNA sequencing and mapping

Small RNA data of *C. grandiflora* and *C. rubella* were obtained from NCBI (Dew-Budd et al., 2024); sRNA sequencing was performed in *C. orientalis* as described before (Dew-Budd et al., 2024). Raw sequencing data from all species was trimmed with TrimGalore v 0.6.5 (Krueger et al., 2021) with parameters --quality 30 --length 19 --max_length 26. The files were filtered to remove reads mapping to the organelle genomes using Bowtie2 (Langmead & Salzberg, 2012). Filtered data was aligned to the respective genomes via Bowtie1 (Langmead et al., 2009) using -q and --best, and converted from SAM to BAM format, then sorted and indexed using SAMtools (Li et al., 2009). Replicates were then merged with the SAMtools merge function. Resulting files were processed using ShortStack v4.1 (Axtell, 2013) and aligned against each genome to get mappability and coverage information. To assess sRNA generation by TEs, BAM files were first filtered for small RNA size with SAMtools view. These were overlapped with the given TE annotation using bedtools –intersect (Quinlan & Hall, 2010).

## Data and Code Availability Statement

Raw data of genome sequencing (PacBio long read seq and OmniC seq) from this article can be found in the NCBI under BioProject numbers: PRJNA1314293 (*C. grandiflora*), PRJNA1314494 (*C. rubella*), and PRJNA1314505 (*C. orientalis*). The *Capsella* genomes and annotations are available on CapBase (www.capsella.uk), figshare (https://doi.org/10.6084/m9.figshare.30048178), and NCBI (*C. rubella* - JBQWLD000000000, *C. orientalis* - JBQWLE000000000, *C. grandiflora hapA* - JBQXEB000000000, *C. grandiflora hapB* - JBQXEC000000000). The ovule RNA-seq data used for gene annotations are available in NCBI via PRJNA1492634. The leaf small RNA-seq data for *Capsellas* can be found via PRJNA1492630 (*C. orientalis*) and PRJNA986946 (*C. rubella* and *C. grandiflora* (Dew-Budd et al., 2024)) in NCBI. The scripts used in this paper are available on GitHub (https://github.com/DrChenHeng/Scripts-to-generate-high-quality-genomic-resource-for-Capsella-species).

## Acknowledgements

This work was supported by the United States National Science Foundation (grant number IOS-2247914 to R.A.M.).

## Author contributions and Conflict of Interest

R.A.M. designed the research; H.C., T.K., and R. A. E. performed research and analyzed data; H.C., R. A. E., and R. A. M. wrote and reviewed the manuscript. The authors declare no competing interests.

---

**Supplementary Table 1.** Generic information of current *Capsella* genome assemblies.

| Species | Taxonomy | Name | ID <sup>a</sup> | Accession | Sequenc<br>e | Annotatio<br>n | Released<br>Year | Source | Reference |
| --- | --- | --- | --- | --- | --- | --- | --- | --- | --- |
| *C. rubella* | 81985 | Capsella rubella v1.1 | GCF_000375325.1 | Monte Gargano | Illumina | Yes | 2013 | Phytozome | Slotte, et al. 2013 |
| *C. rubella* | 81985 | Crubella_RAM_v1.1 | JBQWLD00000000 | Monte Gargano | PacBio & HiC | Yes | 2025 | Mosher Lab | \ |
| *C. grandiflora* | 264402 | Capsella grandiflora v1.1 | Phytozome ID: 266 | 83.17 | Illumina | Yes | 2013 | Phytozome | Slotte et,al. 2013 |
| *C. grandiflora* | 264402 | UAZ_Capgrnd_2 | GCA_040086895.1 | 83.17 | ONT | N/A | 2024 | NCBI | Dew-Budd, et al. 2024 |
| *C. grandiflora* | 264402 | Cgrandiflora_new | GCA_044048995.1 | female (Cg89.3 x Cg81) x male(Cg89.16 x Cg94) | ONT | N/A | 2024 | NCBI | Dziasek, et al. 2024 |
| *C. grandiflora* | 264402 | Cgrandiflora_RAM_v1.1 | JBQXEB00000000 & JBQXEC00000000 | 83.17 | PacBio & HiC | Yes | 2025 | Mosher Lab | \ |
| *C. orientalis* | 945839 | CapOri_SW_1.1 | Genome ID:24034 | Klokov | Illumina | N/A | 2015 | CoGe | Douglas, et al. 2015 |
| *C. orientalis* | 945839 | Corientalis_genome | GCA_964656795.1 | unknown | ONT & HiC | Yes | 2025 | figshare | Kasianova, et al. 2025 |
| *C. orientalis* | 945839 | Corientalis_RAM_v1.1 | JBQWLE00000000 | Klokov | PacBio & HiC | Yes | 2025 | Mosher Lab | \ |

<sup>a</sup> **ID:** The official ID of released assemblies. The IDs of two assemblies, *Capsella grandiflora* v1.1 and CapOri\_SW\_1.1, are from the Phytozome and CoGe database, respectively. IDs for others are from NCBI GeneBank.

**Supplementary Table 2.**
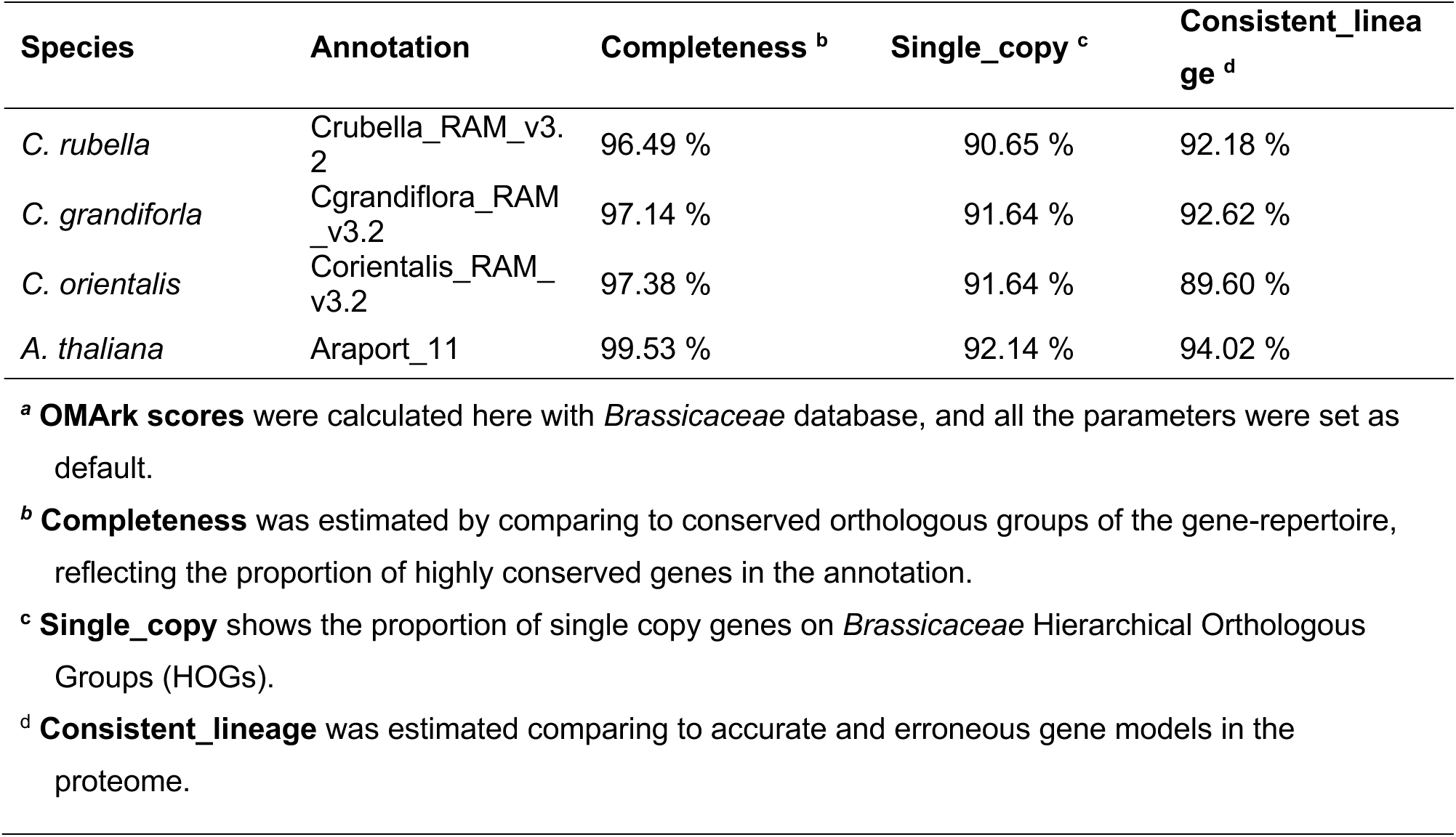
The comparison of *Capsella* to *A. thaliana* genome annotation based on OMArk scores.

| Species | Annotation | Completeness <sup>b</sup> | Single_copy <sup>c</sup> | Consistent_lineage <sup>d</sup> |
| --- | --- | --- | --- | --- |
| *C. rubella* | Crubella_RAM_v3.2 | 96.49 % | 90.65 % | 92.18 % |
| *C. grandiflora* | Cgrandiflora_RAM_v3.2 | 97.14 % | 91.64 % | 92.62 % |
| *C. orientalis* | Corientalis_RAM_v3.2 | 97.38 % | 91.64 % | 89.60 % |
| *A. thaliana* | Araport_11 | 99.53 % | 92.14 % | 94.02 % |

<sup>a</sup> **OMArk scores** were calculated here with *Brassicaceae* database, and all the parameters were set as default.
<sup>b</sup> **Completeness** was estimated by comparing to conserved orthologous groups of the gene-repertoire, reflecting the proportion of highly conserved genes in the annotation.
<sup>c</sup> **Single\_copy** shows the proportion of single copy genes on *Brassicaceae* Hierarchical Orthologous Groups (HOGs).
<sup>d</sup> **Consistent\_lineage** was estimated comparing to accurate and erroneous gene models in the proteome.

**Supplementary Table 6.** The statistics of Capsella transposon annotation at the class and subclass level.

|  | Crubella_RAM_v1 |  |  | Corientalis_RAM_v1 |  |  | Cgrandiflora_RAM_v1_hap A |  |  | Cgrandiflora_RAM_v1_hap B |  |  |
| --- | --- | --- | --- | --- | --- | --- | --- | --- | --- | --- | --- | --- |
| Repeats type | count | length (Mb) | genome (%) | count | length (Mb) | genome (%) | count | length (Mb) | genome (%) | count | length (Mb) | genome (%) |
| **ClassI TE** | 8359 | 9.80 | 5.83 | 9702 | 11.09 | 6.00 | 16433 | 37.86 | 13.60 | 14304 | 18.08 | 7.83 |
| LTR | 6434 | 8.65 | 5.15 | 7685 | 9.99 | 5.41 | 12551 | 35.53 | 12.76 | 10801 | 16.05 | 6.96 |
| SINE | 1045 | 0.23 | 0.14 | 979 | 0.21 | 0.11 | 687 | 0.18 | 0.07 | 2550 | 1.86 | 0.81 |
| LINE | 880 | 0.92 | 0.55 | 1038 | 0.89 | 0.48 | 3195 | 2.15 | 0.77 | 953 | 0.16 | 0.07 |
| **ClassII TE** | 14751 | 10.74 | 6.39 | 17550 | 11.29 | 6.11 | 10585 | 7.77 | 2.79 | 11279 | 8.25 | 3.58 |
| TIR | 1802 | 1.84 | 1.09 | 2539 | 2.26 | 1.22 | 2024 | 1.50 | 0.54 | 2827 | 1.82 | 0.79 |
| MITE | 4875 | 1.52 | 0.91 | 5519 | 1.64 | 0.89 | 829 | 0.19 | 0.07 | 689 | 0.19 | 0.08 |
| nMITE <sup>a</sup> | 3698 | 2.08 | 1.24 | 3865 | 2.46 | 1.33 | 0 | 0.00 | 0.00 | 0 | 0.00 | 0.00 |
| RC | 4376 | 5.30 | 3.16 | 5627 | 4.92 | 2.66 | 7732 | 6.07 | 2.18 | 7763 | 6.24 | 2.71 |
| **Non-TE Repeats** | 9889 | 30.41 | 18.10 | 13416 | 49.42 | 26.75 | 30406 | 92.58 | 33.26 | 16821 | 29.83 | 12.93 |
| tRNA | 0 | 0.00 | 0.00 | 91 | 0.03 | 0.02 | 11 | 0.00 | 0.00 | 11 | 0.00 | 0.00 |
| rRNA | 904 | 0.72 | 0.43 | 33 | 0.61 | 0.33 | 0 | 0.00 | 0.00 | 0 | 0.00 | 0.00 |
| Tandam Repeats | 1102 | 28.65 | 17.05 | 1250 | 46.12 | 24.96 | 7523 | 91.28 | 32.79 | 1060 | 26.84 | 11.63 |
| Satellite | 7261 | 1.51 | 0.90 | 10117 | 2.91 | 1.58 | 22854 | 7.88 | 2.83 | 15750 | 5.47 | 2.37 |
| Simple Repeats | 226 | 0.07 | 0.04 | 216 | 0.07 | 0.04 | 0 | 0.00 | 0.00 | 0 | 0.00 | 0.00 |
| Unknown | 396 | 0.55 | 0.33 | 1709 | 0.86 | 0.47 | 18 | 1.15 | 0.41 | 0 | 0.00 | 0.00 |
| **All TE** | 23110 | 20.54 | 12.22 | 27252 | 22.38 | 12.11 | 27018 | 45.63 | 16.39 | 25583 | 26.33 | 11.41 |
| **Total** | 32999 | 50.95 | 30.32 | 40668 | 71.80 | 38.86 | 57424 | 138.22 | 49.65 | 42404 | 56.16 | 24.34 |

<sup>a</sup> nMITE (not MITE TEs) includes some class II TEs, whose TR domains were identified with length $\geq 800$ bp, which is classified by DeepTE.

**Supplementary Table 7.**
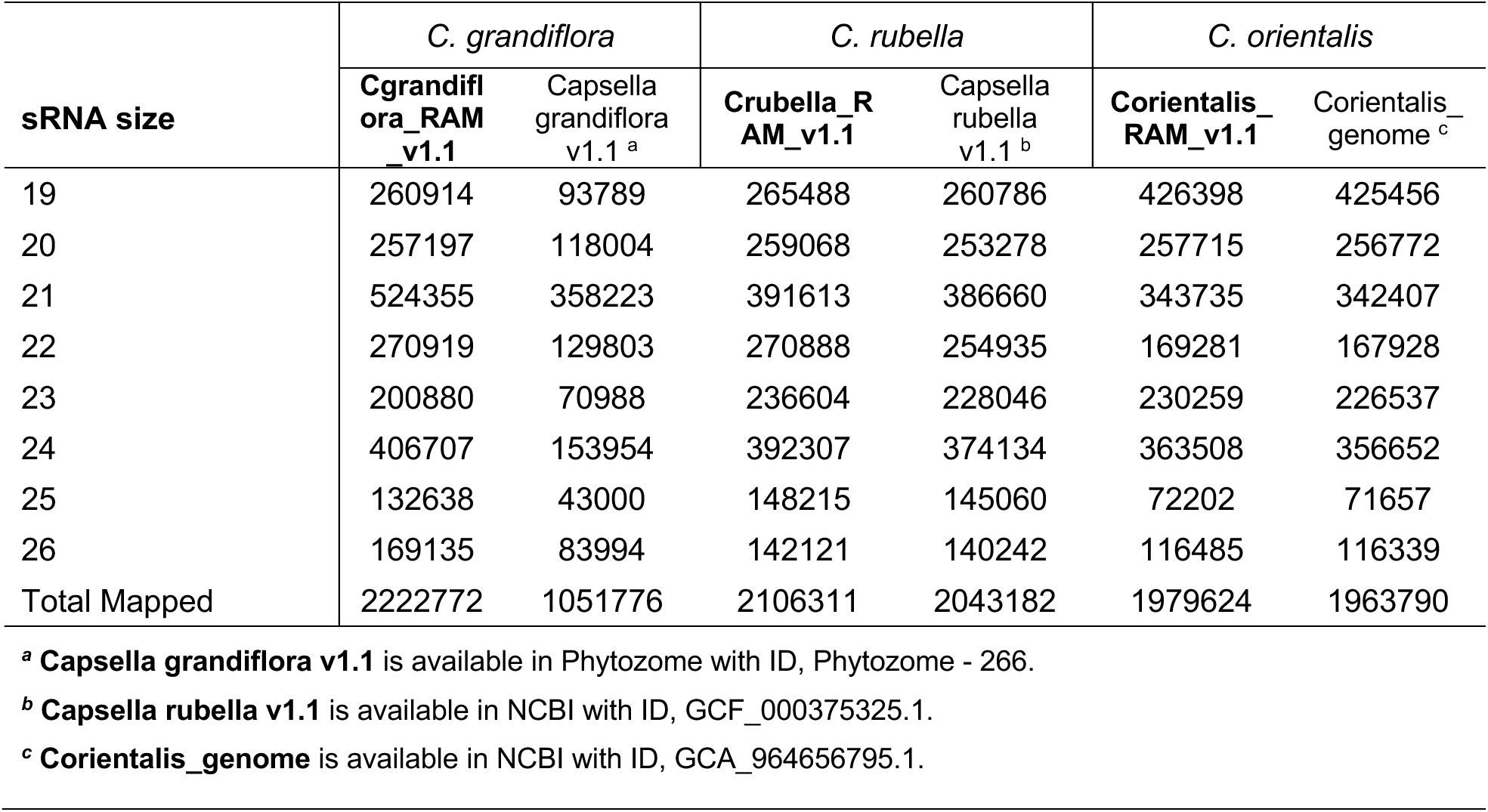
Counts of mapped small RNA over our assemblies versus the publicly available versions.

| sRNA size | *C. grandiflora* |  | *C. rubella* |  | *C. orientalis* |  |
| --- | --- | --- | --- | --- | --- | --- |
|  | **Cgrandiflora_RAM_v1.1** | Capsella grandiflora v1.1 <sup>a</sup> | **Crubella_RAM_v1.1** | Capsella rubella v1.1 <sup>b</sup> | **Corientalis_RAM_v1.1** | Corientalis_genome <sup>c</sup> |
| 19 | 260914 | 93789 | 265488 | 260786 | 426398 | 425456 |
| 20 | 257197 | 118004 | 259068 | 253278 | 257715 | 256772 |
| 21 | 524355 | 358223 | 391613 | 386660 | 343735 | 342407 |
| 22 | 270919 | 129803 | 270888 | 254935 | 169281 | 167928 |
| 23 | 200880 | 70988 | 236604 | 228046 | 230259 | 226537 |
| 24 | 406707 | 153954 | 392307 | 374134 | 363508 | 356652 |
| 25 | 132638 | 43000 | 148215 | 145060 | 72202 | 71657 |
| 26 | 169135 | 83994 | 142121 | 140242 | 116485 | 116339 |
| Total Mapped | 2222772 | 1051776 | 2106311 | 2043182 | 1979624 | 1963790 |

<sup>a</sup> **Capsella grandiflora v1.1** is available in Phytozome with ID, Phytozome - 266.
<sup>b</sup> **Capsella rubella v1.1** is available in NCBI with ID, GCF\_000375325.1.
<sup>c</sup> **Corientalis\_genome** is available in NCBI with ID, GCA\_964656795.1.

**Supplementary Table 8.** Statistics of small RNA mapping for *Capsellas*.

| Data <sup>a</sup> | Raw reads | Filtered reads <sup>b</sup> | Size selected (19-26) | Mapped reads |
| --- | --- | --- | --- | --- |
| *C. grandiflora* rep 1 | 5223465 | 2498098 | 874975 | 810871 |
| *C. grandiflora* rep 2 | 4734947 | 2287822 | 801158 | 753482 |
| *C. grandiflora* rep 3 | 4863134 | 2397153 | 952460 | 888523 |
| *C. rubella* rep 1 | 2841756 | 2047225 | 650782 | 611565 |
| *C. rubella* rep 2 | 4760937 | 2195791 | 772969 | 714890 |
| *C. rubella* rep 3 | 4659037 | 3542215 | 1011979 | 960452 |
| *C. orientalis* rep 1 | 32750007 | 2515468 | 849440 | 799092 |
| *C. orientalis* rep 2 | 30803141 | 1955568 | 582130 | 555169 |
| *C. orientalis* rep 3 | 31682041 | 2659392 | 785687 | 744262 |

<sup>a</sup> **Data** were small RNA seq data from leaves for *C. rubella*, *C. grandiflora*, and *C. orientalis*.
<sup>b</sup> **Filtered reads** count structural small RNAs, like rRNA and tRNA, that are filtered out before mapping.

**Supplementary Figure 1.**
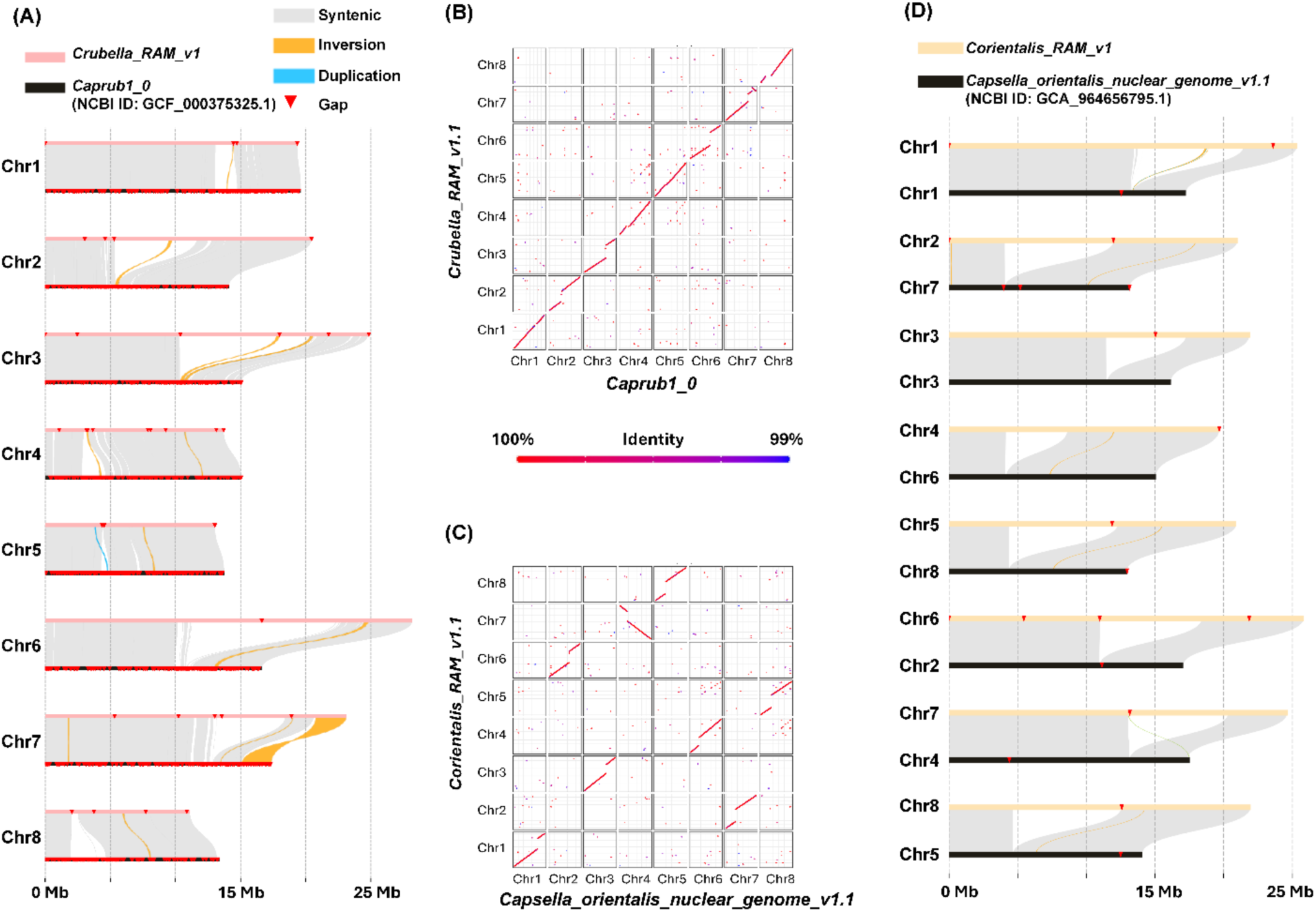
The genome variation among current *Capsella* genomes and public assemblies. (A -. **B)** k-*mer* riparian plot and gene-based synteny map comparing *C. rubella* assemblies. The riparian plot was generated by minimap2 and plotsr, while the gene-based synteny map was generated by customized scripts, see GitHub for more information. **(C - D)** k-*mer* riparian plot and gene-based synteny map comparing *C. orientalis* assemblies. Chr 4 of the public genome (the Chr 7 of the genome Corientalis_RAM_v1) in k-*mer* riparian plot was manually reversed to match the new assembly. Details of each assembly can be found in Table 1 and Table S1.

**Supplementary Figure 2.**
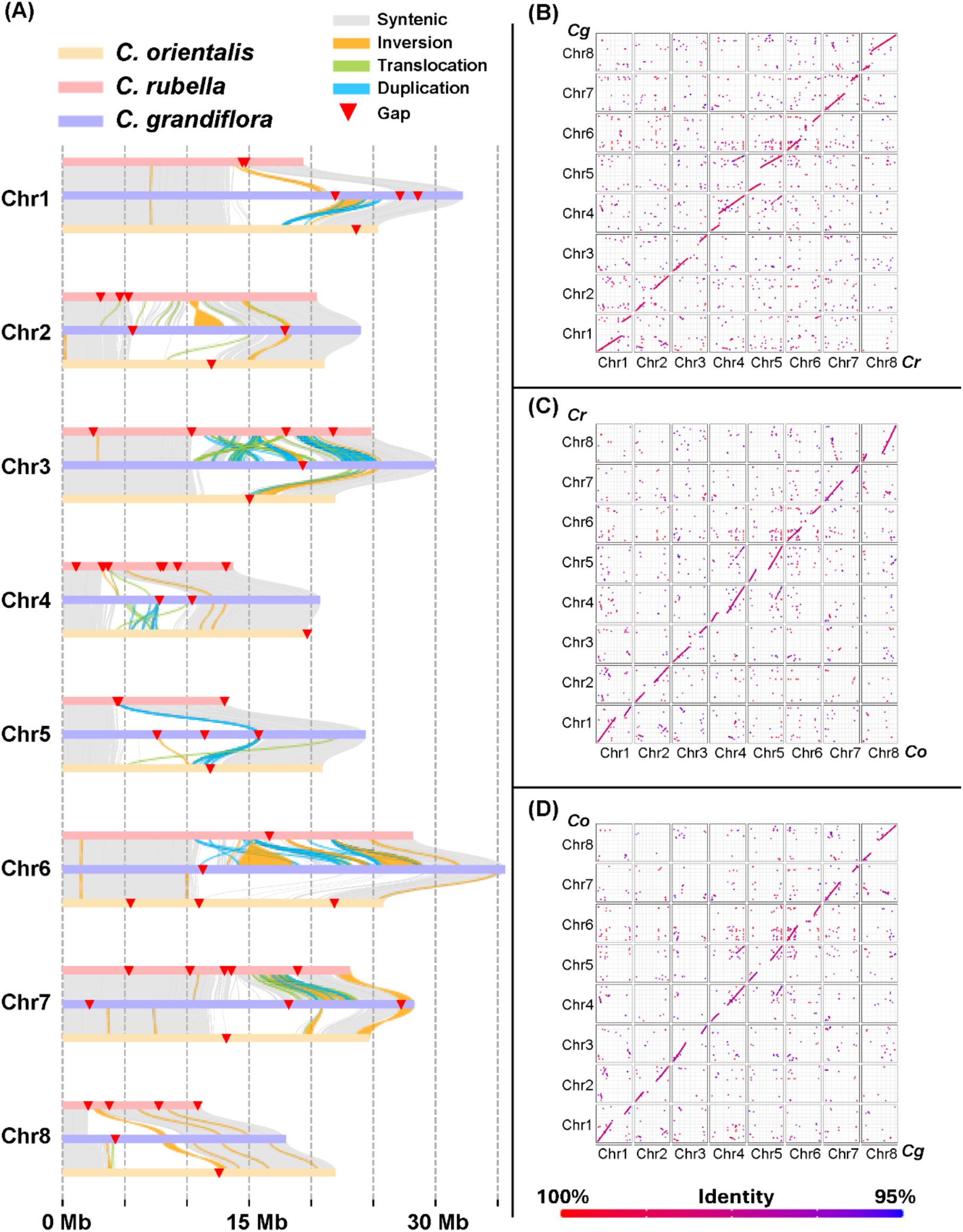
The genome variation among three *Capsella* genomes. **(A)** k-*mer* based riparian plot of three *Capsella* genomes; (B) Gene based synteny map comparing *C. rubella* and *C. grandiflora*; (C) Gene based synteny map comparing *C. orientalis* and *C. rubella*; **(D)** Gene based synteny map comparing *C. grandiflora* and *C. orientalis*.

**Supplementary Figure 3.**
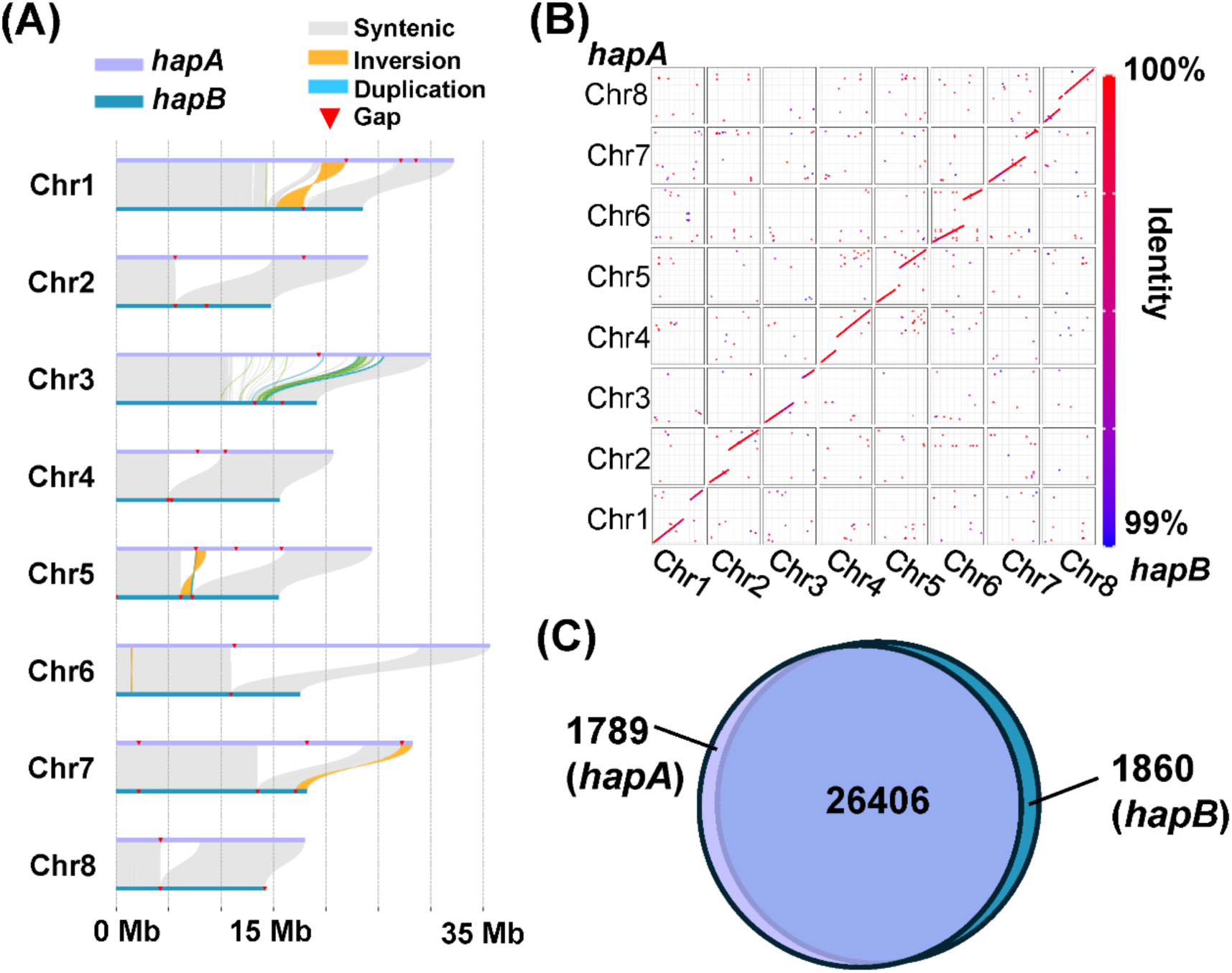
The genome variation between haplotypes of *C. grandiflora*. **(A)** k-*mer* based riparian plot. **(B)** Gene-based synteny map. **(C)** Venn diagram depicting homologous genes annotated in each haplotype. The 26,406 orthologs were defined based on *hapA*. The number slightly varies in *hapB* (26,045), due to haplotype-specific paralogs.

**Supplementary Figure 4.**
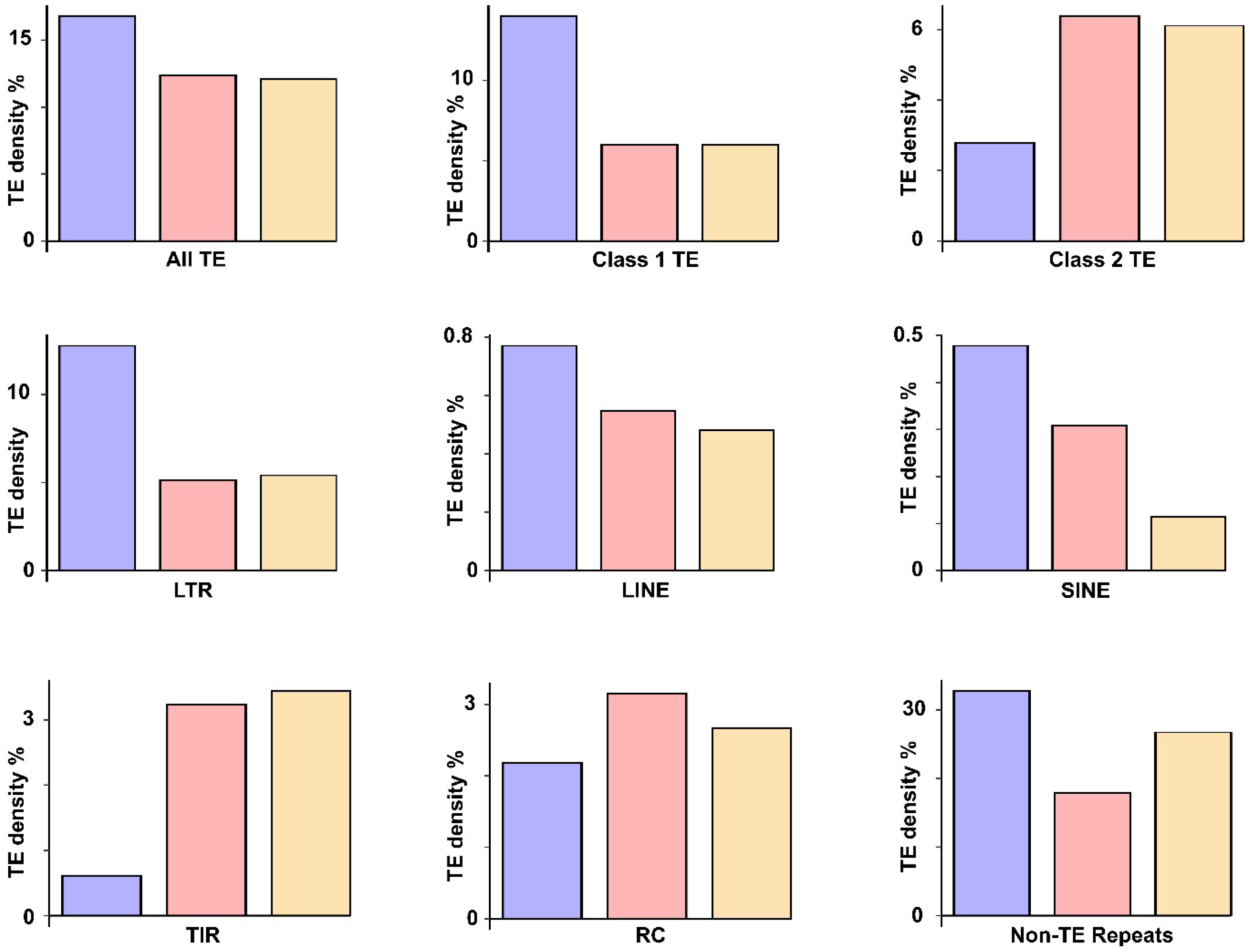
TEs are a larger percentage of the *C. grandiflora* genome compared to *C. rubella* and *C. orientalis*. The TIR includes TIR, MITE and nMITE elements. The non-TE repeats include Satellite DNA, Tandem repeats, Simple repeats, rRNA, tRNA, and Unknown repeats. The purple, pink, and yellow bars are *C. grandiflora*, *C. rubella*, and *C. orientalis*, respectively.

## Online Supplementary Materials

**Supplementary File 1.** All chloroplast and mitochondrial assemblies for *Capsellas*

**Supplementary Table 3**. *C. rubella* genes and their homologies in *Capsella* species and *A. thaliana*

**Supplementary Table 4.** *C. orientalis* genes and their homologies in *Capsella* species and *A. thaliana*

**Supplementary Table 5.** *C. grandiflora* genes and their homologies in *Capsella* species and *A. thaliana*

**Supplementary Table 9.** RNA-seq data for Genome annotation

## Notes

https://figshare.com/articles/dataset/High_quality_genomic_resource_for_i_Capsella_i_species/30048178

https://github.com/DrChenHeng/Scripts_to_generate_high_quality_genomic_resource_for_Capsella_species

### Competing Interest Statement

The authors have declared no competing interest.

### Summary of Updates

Main text re-uploaded to correct Figure 3, which was misformated in first version.

